# Thalamic Nuclei Volumes Across Psychiatric and Neurological Disorders: A Multi-Site Magnetic Resonance Imaging Study

**DOI:** 10.1101/2025.04.21.649628

**Authors:** V. Mäki-Marttunen, S. Nerland, K.N. Jørgensen, E.A. Høgestøl, J. Rokicki, D. Alnæs, S. Borgwardt, B. Boye, J Buitelaar, E. Bøen, S. Cervenka, A. Conzelmann, S. Erhardt, B. Franke, G. Celius E., H.F. Harbo, E. Hilland, P. Hoekstra, C.A. Hartman, R. Jonassen, G. Jönsson E., N.I. Landrø, K-P. Lesch, L.A. Maglanoc, P. Pauli, C.M. Sellgren, G.O. Nygaard, J. Oosterlaan, A. Schmidt, E. Schwarz, G.C. Ziegler, I. Agartz, L.T. Westlye, O.A. Andreassen, T. Kaufmann, T Elvsåshagen

**Author notes:** Corresponding Author: V.Mäki-Marttunen, Centre for Precision Psychiatry, Institute of Clinical Medicine, University of Oslo & Division of Mental Health and Addiction, Oslo University Hospital, Postboks 4956 Nydalen, 0424 Oslo, TE.

## Abstract

The human thalamus is an integrative hub for multiple cortical and subcortical circuits involved in sensory processing and higher cognitive functions. Thalamic volume differences have been reported across multiple psychiatric and neurological disorders, but previous studies have typically relied on small samples, focused on one or a limited number of disorders, or investigated the thalamus as a whole without considering its functional subdivisions. In this multi-site study, we compared thalamic nuclei volumes across mild cognitive impairment (MCI), dementia (DEM), major depressive disorder, schizophrenia spectrum disorder (SCZ), clinical high risk for schizophrenia, bipolar spectrum disorder, autism spectrum disorder, attention-deficit/hyperactivity disorder, Parkinson’s disease, multiple sclerosis (MS), and healthy controls (N > 8 000). Using structural MRI, we segmented 25 bilateral thalamic nuclei, corresponding to six anatomical groups. Linear models revealed that anterior, medial and lateral regions of the thalamus were significantly smaller in several conditions, with largest effects observed for MCI, DEM, SCZ and MS. In contrast, the ventral and intralaminar groups were relatively normal. This pattern of effects largely corresponds to the canonical functional subdivision of the thalamus into higher-order and sensory regions. At the level of individual nuclei, the clinical conditions were associated with distinct patterns of alterations, while left and right lateral geniculate nuclei were implicated in six of the disorders, suggesting a possible relation with circadian and sleep disturbances. Together, the results highlight a role for the higher-order thalamus in common brain disorders and a differential involvement at the nuclei level, refining our understanding of thalamic pathology across common brain disorders.

## Introduction

Common psychiatric and neurological disorders often display comorbidity (1, 2), symptom overlap (3-7), and considerable phenotypic variability within a single disorder (8, 9), posing a need for cross-diagnostic characterizations (10, 11). Large-scale studies integrating samples from multiple sites across clinical conditions have the potential to reveal both shared and unique features and mechanisms. In support of this, recent trans-diagnostic studies revealed substantial genetic overlap and shared structural and functional brain characteristics across a range of disorders (8, 12-19). In this context, the thalamus is of particular relevance, as several of its subregions are implicated in key sensorimotor and cognitive functions that are disrupted in common brain disorders (15, 20). Structural differences in the thalamus have been reported across various psychiatric and neurological disorders (21-28). Previous magnetic resonance imaging (MRI) studies have primarily focused on whole thalamus volumes, most commonly finding decreases in brain disorders. These reductions may result from general loss of thalamic tissue, possibly reflecting illness progression (25, 29). Importantly, measuring whole thalamic volume may have masked subregion-specific effects – for instance, evidence indicates that both increases and decreases in volume of specific thalamic nuclei can occur simultaneously (30, 31). To further understand the involvement of the thalamus in the pathophysiology of brain disorders, the characteristics of thalamic subdivisions warrant further investigation in a cross-diagnostic setting (32, 33).

The human thalamus comprises multiple nuclei that participate in most aspects of brain function, from modulation of arousal and sensory processing to higher cognitive functions (20, 34-38), and jointly play key roles in multiple cortical and subcortical circuits (39-44). Consistent with this functional heterogeneity, projection patterns differ across the thalamus: some regions contain a larger proportion of neurons that relay sensory and motor information to the cortex, such as ventral and geniculate posterior regions, while others contain a larger proportion of neurons interconnected with cortical and limbic regions that participate in cognitive functions, such as the anterior, medio-dorsal and lateral regions (45, 46). The early characterisation of thalamic neurons into thalamic core and matrix based on different patterns of gene expression throughout the thalamus, reflects this functional distinction (47-49). However, to what extent anatomical or functional characterisations are useful for detecting structural changes in psychiatric and neurological disorders remains to be investigated. This is particularly relevant since some thalamic regions can be targeted by neurostimulation interventions (50).

Aiming to discern shared and unique patterns of thalamic abnormalities across neurological and psychiatric disorders, we analysed brain scans from > 8 000 individuals from multiple sites and quantified volumes of 25 thalamic nuclei. We compared nuclei volumes across 11 clinical conditions including mild cognitive impairment (MCI), dementia (DEM), Parkinson’s disease (PD), multiple sclerosis (MS), major depression (MDD), clinical high-risk for schizophrenia (SCZRISK), schizophrenia spectrum disorder (SCZ), bipolar spectrum disorder (BD), non-SCZ psychosis spectrum diagnoses (PSYMIX), autism spectrum disorder (ASD), attention-deficit/hyperactivity disorder (ADHD), and healthy controls (HC). We hypothesized that the clinical conditions would be characterized by distinct patterns of alterations when compared to healthy controls. In addition to case-control comparisons, we investigated the associations between thalamic group volumes and clinical characteristics, i.e., symptoms severity in SCZ and MS and cognitive status in MCI, DEM and HC. To examine potential longitudinal changes in thalamic volumes, we analysed follow-up thalamic volume data available from the MS cohort. Finally, to relate structural alterations to function, we compared the affected thalamic regions with a known functional characterization, the higher-order thalamic matrix and the thalamic sensory core, based on mRNA expression. Our study highlights the heterogeneity of thalamic involvement across psychiatric and neurological disorders.

## Methods

### Samples and thalamus segmentation

We obtained data from 9 646 individuals (n = 5 094 patients, n = 4 552 HC) through collaborations, data sharing platforms, and in-house samples (Table S1). The patient samples included individuals with ADHD, ASD, BD (type I and II), DEM, MCI, MDD, MS, PD, SCZ (including individuals with schizoaffective disorder), SCZRISK and PSYMIX. DEM and MCI were defined based on a battery of tests (adni.loni.usc.edu/data-samples/adni-data/study-cohort-information, and Ref. (51)). Some individuals with BD presented psychotic symptoms. Data collection for each sample was performed with the written informed consent of participants and with approval by the respective local Institutional Review Boards.

For the segmentation of the thalamus, we used T1-weighted three-dimensional brain MRI data. Using Bayesian thalamus segmentation based on ex-vivo MRI and histology in Freesurfer 6.0 (52), we segmented the MRI data into the whole thalamus and 25 bilateral thalamic nuclei (Figure 1). Following previous work, including our own (15,41), we assigned the thalamic nuclei to six thalamic anatomical groups (Table 1). The thalamic segmentation has high test–retest reliability (most nuclei having intra-class correlation coefficient above 0.90) and is robust to differences in MRI contrast (26, 52). It also shows good agreement with histological divisions of thalamic nuclei (52), and the six groups have been used in a previous multi-diagnostic study (15). To ensure data quality, V.M.M visually inspected all images and corresponding segmentations in the MRI datasets by examining axial and coronal view figures of the segmentations for each participant (see Supplementary material). Volumes with low image quality, artifacts, or failed thalamic segmentations were excluded, yielding the final samples described in Table 2. For the main analyses, we extracted volumetric measures in mm^3^ for each thalamic group (combining left and right sides), and used these data for statistical analyses.

**Table 1.** Thalamic groups definition and volume.

| Group | Nuclei | Reference volume (mm<sup>3</sup>) |
| --- | --- | --- |
| **Anterior** | anteroventral | 261 |
| **Lateral** | laterodorsal and lateral posterior | 297 |
| **Medial** | paratenial, reuniens, mediodorsal medial magnocellular, and mediodorsal lateral parvocellular nucleus | 2221 |
| **Ventral** | ventral anterior, ventral anterior magnocellular, ventral lateral anterior, ventral lateral posterior, ventral posterolateral, and ventromedial | 5454 |
| **Intralaminar** | central medial, central lateral, paracentral, centromedial, and parafascicular nucleus | 807 |
| **Posterior** | lateral geniculate, medial geniculate, limitans, pulvinar anterior, pulvinar medial, pulvinar lateral, and pulvinar inferior | 4262 |

**Table 2.**
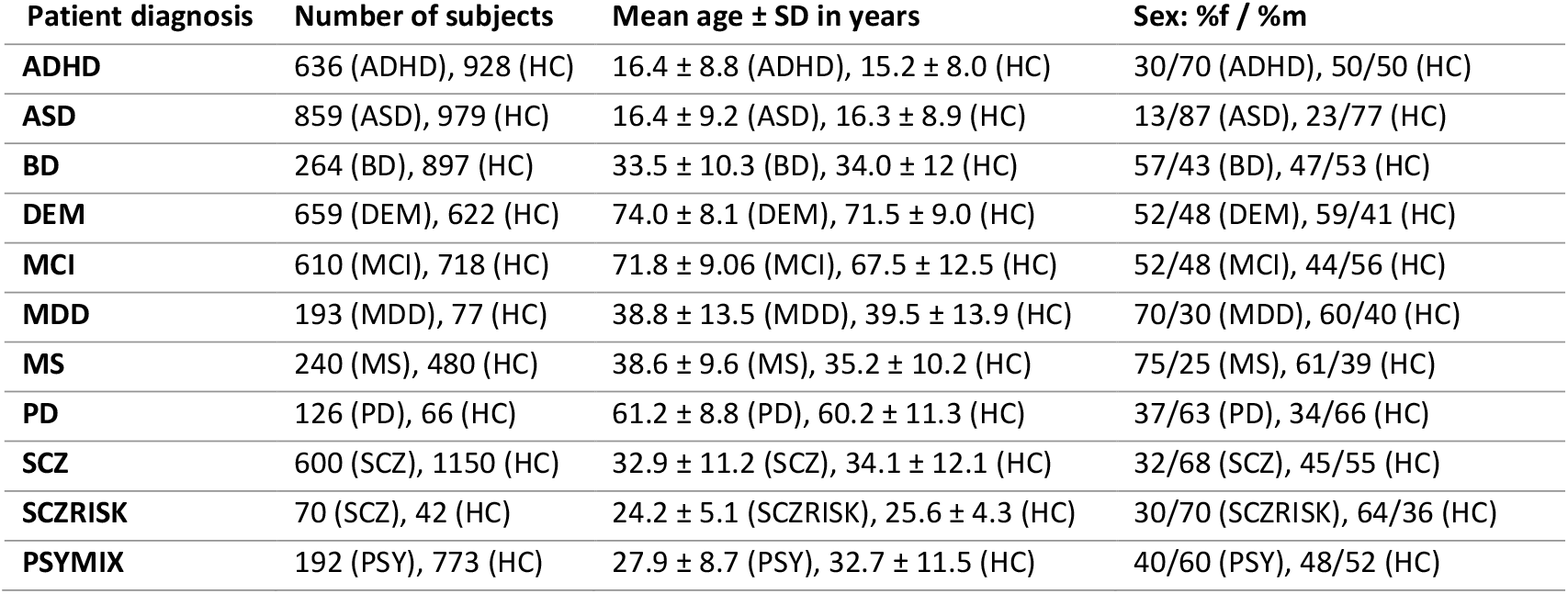
Size and demographic characteristics of the samples, grouped by patient diagnosis. ADHD: attention deficit/hyperactivity disorder; ASD: autism spectrum disorder; BD: bipolar disorder; DEM: dementia; MCI: mild cognitive impairment; MDD: major depressive disorder; MS: multiple sclerosis; PD: Parkinson’s disease; SCZ: schizophrenia; SCZRISK: individuals at clinical high-risk for schizophrenia; PSYMIX: non-SCZ psychosis spectrum diagnoses; HC: healthy controls.

| Patient diagnosis | Number of subjects | Mean age ± SD in years | Sex: %f / %m |
| --- | --- | --- | --- |
| **ADHD** | 636 (ADHD), 928 (HC) | 16.4 ± 8.8 (ADHD), 15.2 ± 8.0 (HC) | 30/70 (ADHD), 50/50 (HC) |
| **ASD** | 859 (ASD), 979 (HC) | 16.4 ± 9.2 (ASD), 16.3 ± 8.9 (HC) | 13/87 (ASD), 23/77 (HC) |
| **BD** | 264 (BD), 897 (HC) | 33.5 ± 10.3 (BD), 34.0 ± 12 (HC) | 57/43 (BD), 47/53 (HC) |
| **DEM** | 659 (DEM), 622 (HC) | 74.0 ± 8.1 (DEM), 71.5 ± 9.0 (HC) | 52/48 (DEM), 59/41 (HC) |
| **MCI** | 610 (MCI), 718 (HC) | 71.8 ± 9.06 (MCI), 67.5 ± 12.5 (HC) | 52/48 (MCI), 44/56 (HC) |
| **MDD** | 193 (MDD), 77 (HC) | 38.8 ± 13.5 (MDD), 39.5 ± 13.9 (HC) | 70/30 (MDD), 60/40 (HC) |
| **MS** | 240 (MS), 480 (HC) | 38.6 ± 9.6 (MS), 35.2 ± 10.2 (HC) | 75/25 (MS), 61/39 (HC) |
| **PD** | 126 (PD), 66 (HC) | 61.2 ± 8.8 (PD), 60.2 ± 11.3 (HC) | 37/63 (PD), 34/66 (HC) |
| **SCZ** | 600 (SCZ), 1150 (HC) | 32.9 ± 11.2 (SCZ), 34.1 ± 12.1 (HC) | 32/68 (SCZ), 45/55 (HC) |
| **SCZRISK** | 70 (SCZ), 42 (HC) | 24.2 ± 5.1 (SCZRISK), 25.6 ± 4.3 (HC) | 30/70 (SCZRISK), 64/36 (HC) |
| **PSY MIX** | 192 (PSY), 773 (HC) | 27.9 ± 8.7 (PSY), 32.7 ± 11.5 (HC) | 40/60 (PSY), 48/52 (HC) |

**Figure 1.**
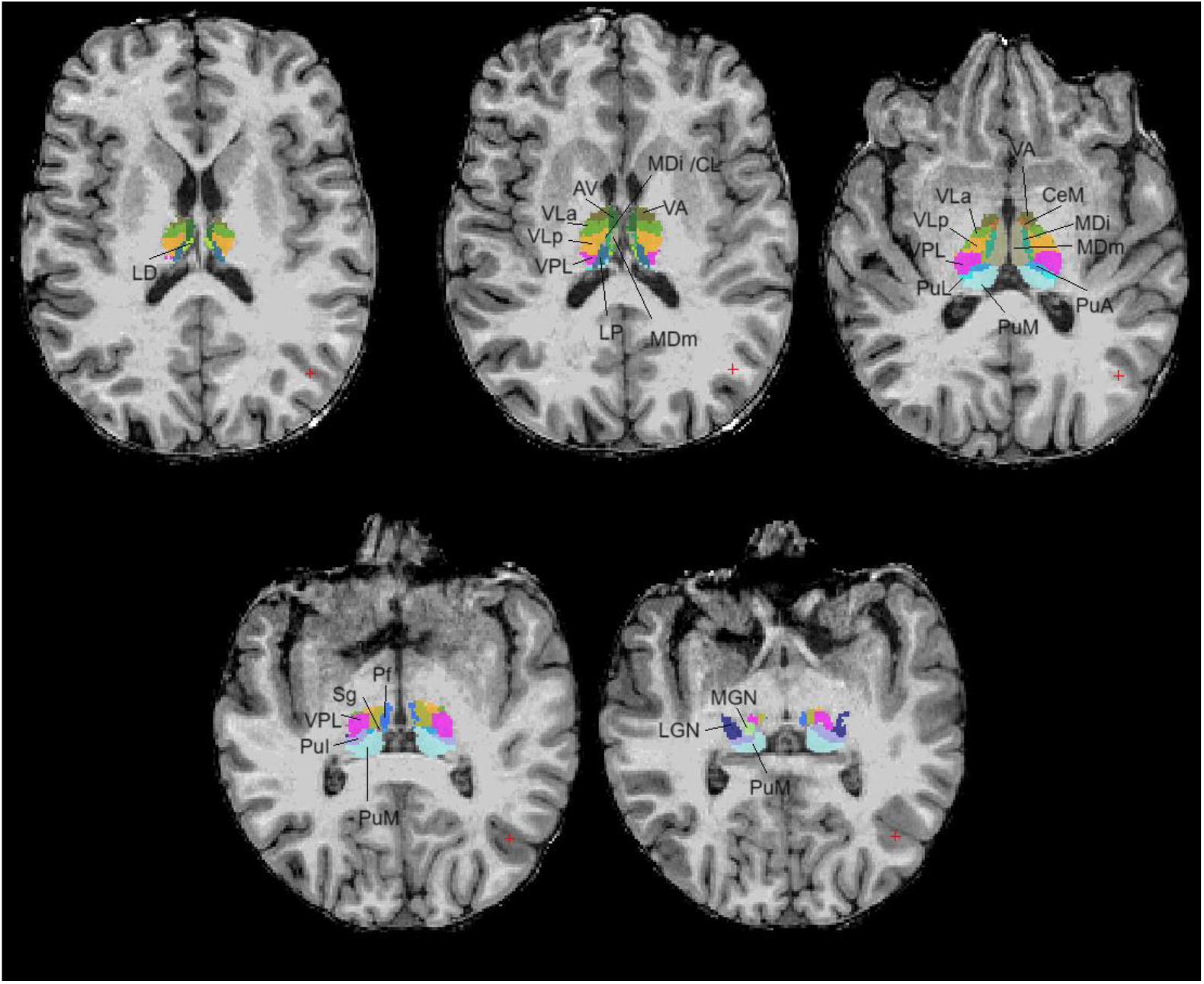
Example of the segmentation of the thalamus of one individual into 50 different nuclei (25 for each hemisphere). LD: latero-dorsal; AV: anteroventral; VLa: ventral lateral anterior; VLp: ventral lateral posterior; VPL: ventral posterolateral; LP: lateral posterior; MDm: medio-dorsal magnocellular; VA: ventral anterior; MDl: mediodorsal lateral; CeM: centro-median; PuA: pulvinar anterior; PuM: pulvinar medial; PuL: pulvinar lateral; PuI: pulvinar inferior; Sg: suprageniculate; Pf: parafascicular; MGN: medial geniculate nucleus; LGN: lateral geniculate nucleus.

### Statistical analysis of volumes, brain disorders, and clinical variables

Statistical analyses for group comparisons were conducted using linear regression models in R v4.2.1 (R Core Team, 2022). Given that thalamic volume changes over the life span (Sullivan et al. 2004), for each diagnosis we matched healthy controls and patients with respect to mean age and age range. All patient samples had controls scanned on the same scanner using identical MRI sequence parameters. For clinical conditions where participants were scanned on multiple scanners, we applied a multivariate scanner harmonization method (**RE**moval of **L**atent **I**nter-scanner **E**ffects through **F**actorization, RELIEF, (53)) that removes scanner-related effects while retaining associations of interest. Unlike other common methods, such as ComBat, which remove explicit scanner effects (e.g. additive and multiplicative), RELIEF also accounts for latent scanner effects (54). As an example, Figure S1 shows the decreasing trend of thalamic volume in healthy controls as a function of age for the anterior and posterior groups before and after scanner harmonization.

For each of the clinical conditions, we fitted linear regression models to test for diagnostic group effects on whole thalamic volume as well as on the volumes of the six thalamic groups. Assumptions of homoscedasticity were met in all cases, except for MS where variances were calculated separately for patients and controls. Volumes were included as dependent variables and diagnosis was the independent variable along with sex, age, age^2^ and estimated intracranial volume (ICV, (55)) as covariates (14). The resulting p-values were adjusted for multiple testing (i.e., 6 comparisons, one per thalamic group) using Bonferroni correction. To test for effects that were independent of global thalamic effects, we performed further analyses including whole thalamic volume as an additional covariate. Effect sizes were calculated using the cohens_d function of the R package effectsize.

To further compare the diagnostic groups, we computed the Pearson’s correlation matrix between the vectors of effect sizes from the case-control comparisons of thalamic groups (Figure 3) (16). Based on this matrix, we derived a distance matrix using the Euclidean distance metric, where higher correlation will result in a smaller distance. On these distances, we performed hierarchical clustering with the function hclust in R using the complete linkage method. The resulting dendrogram positions closer groups together, with bar height representing the distance (Figure 3).

To explore differences at the level of individual nuclei, we conducted linear regression models on the data harmonized within each patient group to test for diagnostic group effects on each nucleus, and included sex, age, age^2^, ICV, and whole thalamic volume as covariates. We adjusted the statistical threshold using Bonferroni correction (i.e., correcting for 50 comparisons, one per nucleus).

Information concerning illness severity was available for individuals with MCI, DEM, MS, SCZ, and PD (Table 3). Linear regression models were fitted to examine relationships between clinical variables (i.e., predictors) and thalamic group volumes (i.e., dependent variables), including also ICV, age, age^2^ and sex.

**Table 3.** Data on clinical assessment and sample sizes. MMSE: Mini-Mental State Examination (56); EDSS: Expanded Disability Status Scale (57); PANSS: Positive and Negative Syndrome Scale (58).

| Sample | Test | Diagnosis | N |
| --- | --- | --- | --- |
| MCI, DEM | MMSE | MCI | 1250 |
| MCI, DEM | MMSE | DEM | 357 |
| MCI, DEM | MMSE | HC | 985 |
| MS | EDSS | MS | 221 |
| SCZ | PANSS | SCA | 200 |

While the cross-sectional analysis on the MS sample was conducted using baseline data from the first scanning instance, MRI and clinical follow-up data was also available from up to six additional time points and was employed to investigate the clinical relevance of longitudinal changes in thalamic volumes (Table S2). For this analysis, scanner-harmonized data corrected for ICV, age, age^2^ (at first session) and sex were entered into a linear mixed model using lme4 package (59). We included volume as the dependent variable and years after first session as the variable of interest, and a random effect was included to account for individual differences.

### RNA Expression analysis

As a comparison for the cross-sectional results, we examined the expression of the thalamic matrix marker calbindin-2 and the core thalamus marker parvalbumin in humans. RNA expression of these proteins was obtained from the available nuclei at the Human Protein Atlas (www.proteinatlas.org). Expression data of all available nuclei within a group (Table S3, Figure S2 and S3) were averaged. As an independent replication, we obtained RNA expression of calbindin-2 and parvalbumin from the Allen Brain Institute (human.brain-map.org, Figure S4). Based on these data, we aggregated the thalamic groups into core thalamus (ventral, intralaminar and posterior groups) and matrix thalamus (lateral and medial groups). We then calculated Cohen’s d effect size for each functional division between the patient and control groups. This allowed us to assess whether the effect sizes differed between core and matrix thalamus by examining the overlap of their confidence intervals.

### Uniform Manifold Approximation and Projection (UMAP) analysis

As a final trans-diagnostic analysis, we aimed to provide a low-dimensional description of the shared and unique patterns of thalamic nuclei volume differences for each diagnostic group relative to healthy controls. To this end, we implemented a dimensionality reduction technique to synthesize the data of thalamic nuclei volumes (i.e. 50 nuclei across left and right hemispheres) across disorders. We first harmonized the data across scanners, including correction for relevant covariates (sex, age, age^2^, ICV) and whole thalamic volume. We then related the volume of each nucleus for each patient to the corresponding average from the sample’s healthy control group, thereby obtaining a measure of effect size. Finally, we projected the data onto a low-dimensional space (2-D) using an unsupervised manifold learning algorithm based on topological analysis, the Uniform Manifold Approximation and Projection (UMAP). This analysis was done with the umap toolbox implemented in Python 3.7 (60, 61). The UMAP approach preserves the global data structure and the local neighbour relations better than other existing methods such as Principal Components Analysis, and has been previously applied to biological data (62, 63). We then visualized the density distributions of individuals with the different diagnoses in the 2-D space. Figure S5 summarizes the pipeline of the UMAP analysis.

## Results

### Case-control comparison of thalamic group volumes

We found significantly larger total thalamic volume in PD, ASD and PSYMIX, with no significant differences in specific thalamic groups. In contrast, BD, SCZ and MS showed significantly reduced whole thalamic volume. Figure 2.a-b and Figure 3.a depict the effect sizes and significant case-control differences for each clinical condition. SCZ, DEM, MCI, and MS showed reduced volume in the lateral and medial thalamic groups. In addition, DEM, MCI and MS exhibited reduced anterior thalamic volume. The reduction in lateral and medial thalamic groups observed in SCZ, DEM, MCI and MS persisted after adjustment for whole thalamic volume, indicating a regional effect beyond the overall thalamic volume decrease seen in e.g. SCZ and MS. The relatively larger ventral and intralaminar thalamic groups found in SCZ, DEM, MCI and MS after correction for whole thalamic volume largely disappeared without such correction, indicating a relative normal volume in these regions with respect to global thalamic effects, rather than an absolute volume increase in these patient groups. We found no significant case-control differences for MDD or SCZRISK. In sum, among the disorders showing specific thalamic groups effects, volume reductions in the anterior, lateral and medial thalamus were most prominent, and the patterns of thalamic group differences effectively clustered the diagnoses with more similar patterns (Figure 3.b and c), demonstrating that thalamic group volumes effectively capture similarities across conditions.

**Figure 2.**
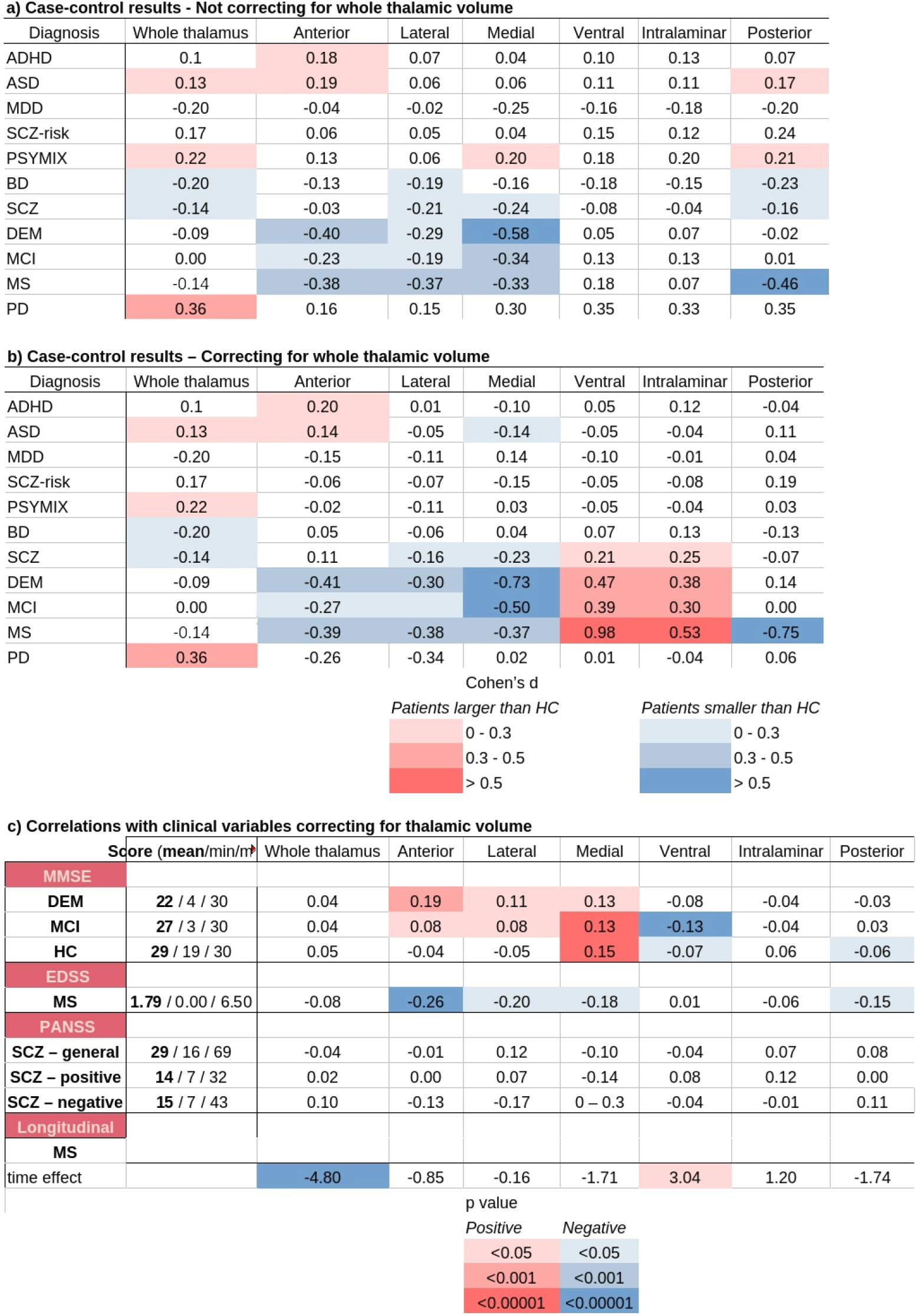
a) Thalamic volumetric differences between patients and healthy controls. Numbers indicate Cohen’s d for the patients versus HC analysis. Only significant comparisons between patients and control groups after correction for multiple comparisons are coloured. Data corrected for scanning site, total ICV, age, age^2^ and sex. b) Same as previous but correcting for whole thalamic volume. c) Coefficients of linear regression between thalamic groups volume and clinical variables. Cells coloured in red indicate significant positive associations and cells in blue indicate significant negative associations. ADHD: attention deficit/hyperactivity disorder; ASD: autism spectrum disorder; BD: bipolar disorder; DEM: dementia; MCI: mild cognitive impairment; MDD: major depressive disorder; MS: multiple sclerosis; PD: Parkinson’s disease; SCZ: schizophrenia; SCZRISK: individuals at clinical high-risk for schizophrenia; PSYMIX: non-SCZ psychosis spectrum diagnoses; HC: healthy controls; MMSE: Mini-Mental State Examination; EDSS: Extended Disability Status Scale; PANSS: Positive and Negative Syndrome Scale.

**Figure 3.**
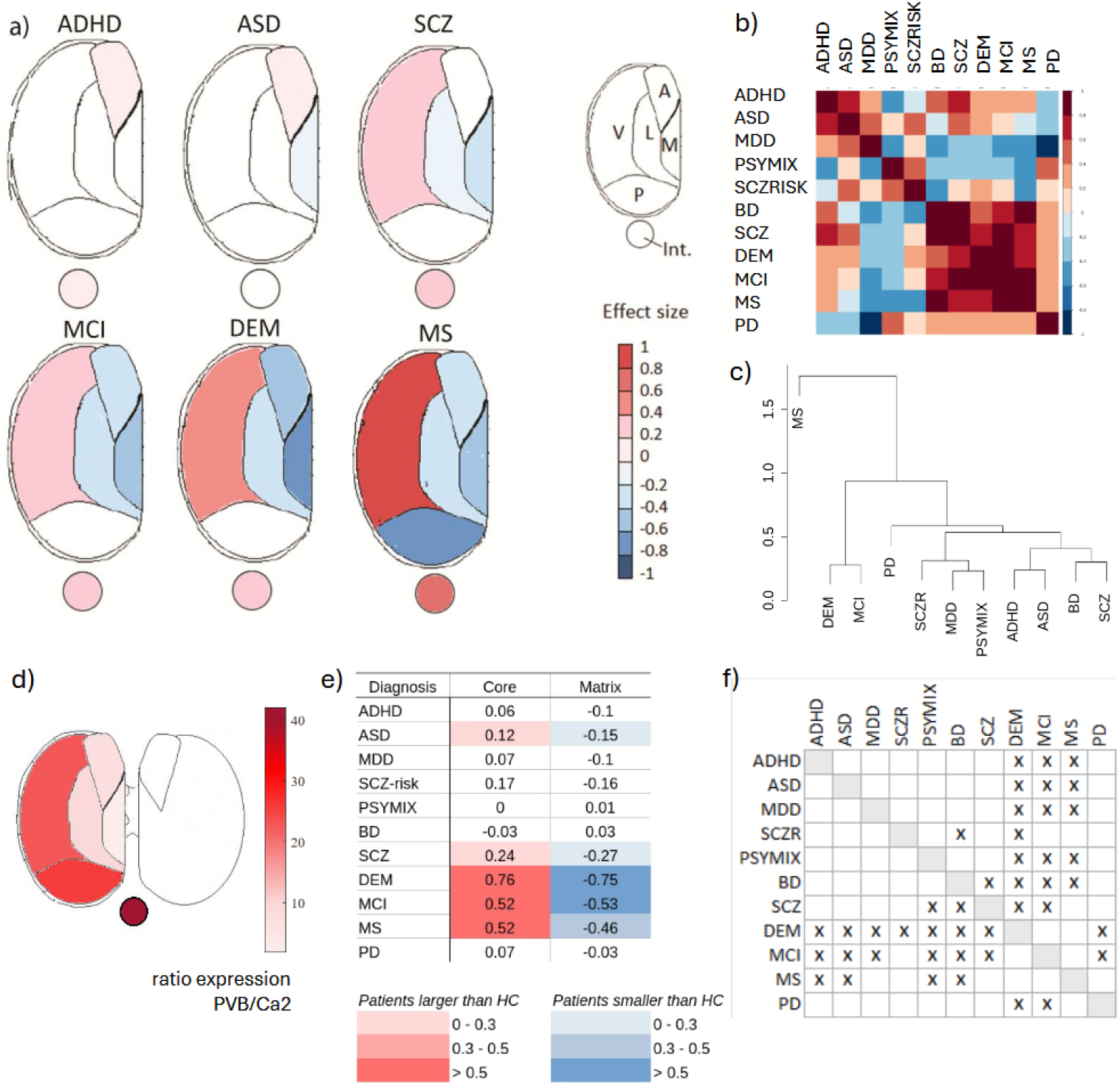
a) Illustration of thalamic groups in the axial plane presenting significant volumetric differences between patients and controls after correction for whole thalamic volume, as shown in Figure 2.b. Results were calculated collapsing the data for left and right hemispheres, and thus only one side is shown. Color bar indicates Cohen’s d. b) Correlation matrix of thalamic group effects across disorders. Larger correlations indicate more similar patterns of case-control differences in thalamic groups (rows in Figure 2.b). Color bar indicates correlation value. For a similar correlation approach at the level of individual nuclei, see Figure S4. c) Clustering dendrogram of the different diagnoses based on the distances obtained from the correlation matrix in b). d) mRNA expression ratio between the core thalamus marker parvalbumin and the thalamic matrix marker calbindin-2 in humans. The colour code indicates the ratio. See also Figure S3 for the expression distribution, and Figure S4 for the expression distribution estimated from an independent dataset (Allen Brain Institute). e) Effect size of difference in thalamic nuclei volumes between patients and controls based on a core-matrix characterization. Coloured cells indicate instances where the effect size confidence intervals of core and matrix groups do not overlap. f) Comparison of effect size confidence intervals between the different diagnoses based on the comparisons in e). Crosses indicate non-overlapping intervals. Upper triangle: core; lower triangle: matrix. ADHD: attention deficit/hyperactivity disorder; ASD: autism spectrum disorder; BD: bipolar disorder; DEM: dementia; MCI: mild cognitive impairment; MDD: major depressive disorder; MS: multiple sclerosis; PD: Parkinson’s disease; SCZ: schizophrenia; SCZRISK: individuals at clinical high-risk for schizophrenia; PSYMIX: non-SCZ psychosis spectrum diagnoses; Ca: calbindin-2; PVB: parvalbumin; A: anterior; V: ventral; L: lateral; M: medial; P: posterior; Int.: intralaminar.

We then compared the pattern of thalamic group alterations with a known functional characterisation of the thalamus into thalamic core and thalamic matrix based on the average expression of the molecular markers calbindin-2 and parvalbumin genes (Figure 3.d and Figures S3 and S4). Ventral, intralaminar and posterior groups showed higher expression of parvalbumin, whereas medial and lateral groups exhibited higher expression of calbindin-2 gene. The anterior group had similarly lower expression of both markers. Based on this characterization, the nuclei groups displaying smaller volumes in patients corresponded primarily to the matrix thalamus. To compare these effects across diagnoses, we aggregated the thalamic group volumes according to the core and matrix subdivisions and calculated the effect size between volumes of patients and controls (Figure 3.e). Comparing the confidence intervals of these effect sizes (Figure 3.f), we observed that the opposite effects for the core and matrix thalamus largely co-occurred. DEM, MCI and MS showed larger effect sizes than the other groups, suggesting the regional segregation between thalamic matrix and core markers may underlie volume abnormalities across different patient groups.

### Case-control comparison of individual thalamic nuclei volume

To further characterize disease-related thalamic volume changes at higher anatomical resolution, we performed group analyses for the individual nuclei volumes and found distinct patterns of case-control differences for the different clinical conditions (Figure 4.a). MS, MCI and DEM presented the largest number of nuclei with significant case-control differences (Figure 4.b), with a high degree of correspondence between the left and right thalamus. Bilateral LGN was the most frequently represented nuclei across disorders (Figure 4.a and c), showing significant case-control differences in five of the 11 conditions. Many of the affected nuclei belonged to the medial group, whereas the posterior group was less frequently involved (Figure 4.a and c).

**Figure 4.**
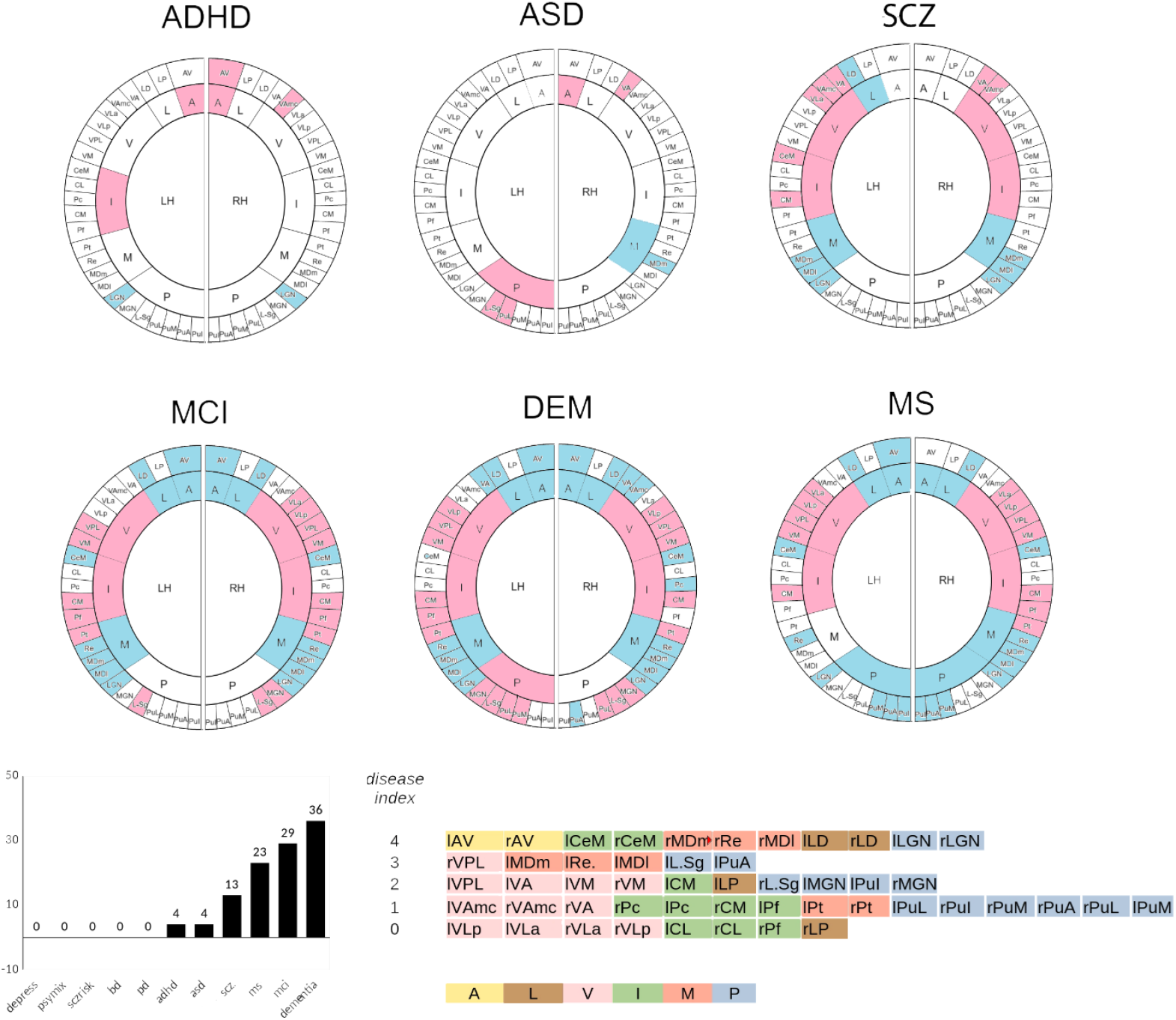
a) Individual thalamic nuclei and lateralized thalamic groups presenting significant case-control differences after correcting for whole thalamic volume and adjusting for relevant covariates. Cells in red correspond to larger volume in patients, while blue corresponds to smaller volume in patients. b) Percentage of individual nuclei presenting significant case-controls differences for the different clinical conditions. c) Number of clinical conditions showing a significant case-control difference (“disease index”) after Bonferroni correction. Zero indicates nuclei with no significant differences. Colours indicate thalamic groups. ADHD: attention deficit and hyperactivity disorder; ASD: autism spectrum disorder; BD: bipolar disorder; DEM: dementia; MCI: mild cognitive impairment; MDD: major depressive disorder; MS: multiple sclerosis; PD: Parkinson’s disease; SCZ: schizophrenia; SCZRISK: individuals at clinical high-risk for schizophrenia; PSYMIX: non-SCZ psychosis spectrum diagnoses; LD: latero-dorsal; AV: anteroventral; VAmc: ventral anterior magnocellular; VLa: ventral lateral anterior; VLp: ventral lateral posterior; VPL: ventral posterolateral; LP: lateral posterior; MDm: medio-dorsal magnocellular; VA: ventral anterior; VM: ventromedial; CL: central lateral; Pc: paracentral; Pt: paratenial; MDl: mediodorsal lateral; CeM: centro-median; PuA: pulvinar anterior; PuM: pulvinar medial; PuL: pulvinar lateral; PuI: pulvinar inferior; Sg: suprageniculate; Pf: parafascicular; MGN: medial geniculate nucleus; LGN: lateral geniculate nucleus; LH: left hemisphere; RH: right hemisphere.

### Association between thalamic volume and clinical measures

Models testing for associations between thalamic group volumes and clinical characteristics (Figure 2.c and Table S4) revealed a significant relation between MMSE scores and thalamic group volumes in DEM and MCI, and in HC. Higher cognitive capacity corresponded to larger volumes, except for the ventral and posterior groups in healthy controls and MCI, where the correlations were negative. The positive correlation between volume and MMSE was found in anterior, lateral and medial groups. We also found a significant correlation between volume of several thalamic groups and EDSS (disability scores) in MS, corresponding to anterior, lateral, medial and posterior groups. Thalamic group volumes did not show significant association with PANSS scores in patients with SCZ.

Using longitudinal data from the participants with MS, we found significant volume decreases across thalamic nuclei volumes and of the whole thalamus, except for the ventral nuclei group (Figure 2.c). The change in volume of the whole thalamus and thalamic groups in follow-up measurements (with respect to baseline) was not significantly correlated with changes in EDSS scores between sessions (p > 0.2).

### Low-dimensional representation of thalamic nuclei volume across disorders

To examine whether collective patterns of thalamic volume alterations emerged when considering all nuclei jointly -i.e. patterns not evident when looking at the different nuclei separately - we implemented a dimensionality reduction technique (UMAP). The diagnoses groups presented distinct patterns of density in the lower-dimensional space (Figure 5, see Figure S7 for annotated density). For instance, patients with MS, DEM, MCI and PD were more prevalent at larger values of the first main axis but differed in the position with respect to the second axis. DEM and MCI presented higher densities (∼38% and ∼27%, respectively) at higher values of the first axis and around zero values of the second axis. MS presented higher density of occurrences (48%) in the upper right quadrant of the 2-D space. Some conditions, e.g. SCZ, presented a wider distribution. Importantly, the distribution of healthy older adults in the 2-D space was markedly different from that of DEM, MCI, and PD – groups with similarly older populations – indicating that age did not drive the observed distribution of groups (Figure S7). Whether the identified clusters (that is, groups with common morphological characteristics, Figure S8) constitute clinically relevant biotypes needs to be explored further. Altogether, the results suggest that the different clinical conditions display differences in the patterns of thalamic nuclei volumetry when considering all nuclei together.

**Figure 5.**
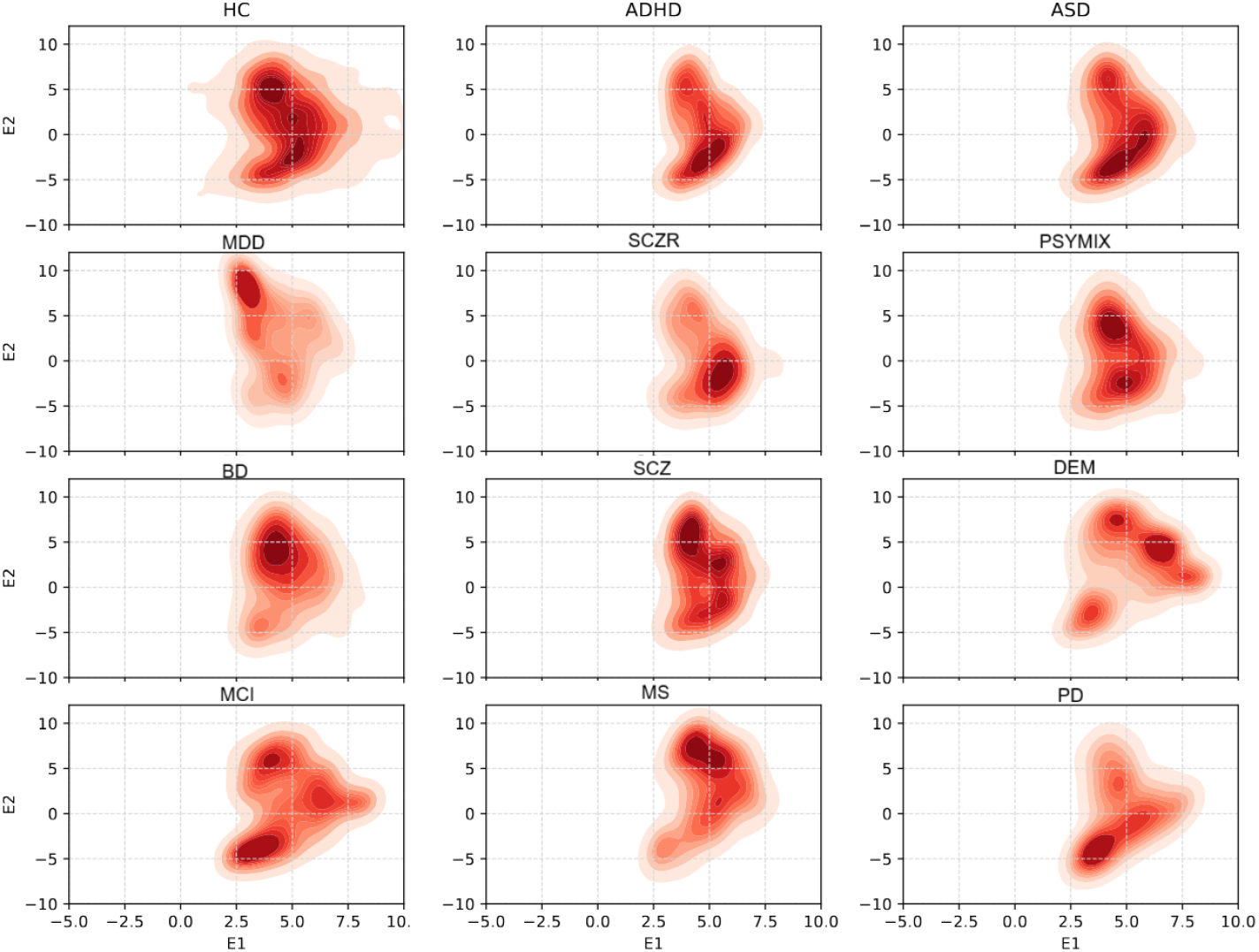
Embedding of all brain disorders in the 2-components space after UMAP dimensionality reduction of the individual thalamic nuclei effects. Colour scale and contours represent the density of occurrences in each area of the 2-D space. Data from each individual for the clinical groups was normalized to the average sample’s healthy control group and were submitted to the UMAP algorithm. ADHD: attention deficit and hyperactivity disorder; ASD: autism spectrum disorder; BD: bipolar disorder; DEM: dementia; MCI: mild cognitive impairment; MDD: major depressive disorder; MS: multiple sclerosis; PD: Parkinson’s disease; SCZ: schizophrenia; SCZRISK: individuals at clinical high-risk for schizophrenia; PSYMIX: non-SCZ psychosis spectrum diagnoses.

## Discussion

In the present study, we analysed 25 bilateral thalamic nuclei volumes in a large dataset comprising patients diagnosed with 11 psychiatric and neurological conditions. SCZ, DEM, MCI and MS presented significant case-control differences across several thalamic groups. Notably, these differences followed a topological distribution across disorders, with smaller volumes in anterior, lateral and medial groups, and larger volumes in ventral and intralaminar groups. Further analyses of individual nuclei, including an embedding in a low dimensional space, revealed differences between clinical conditions. Our results show disease-specific patterns of thalamic volumes and general brain disorder patterns across clinical conditions. These findings point both to commonalities and differences across clinical conditions, contributing to a more detailed characterisation of alterations in the thalamus across brain disorders.

### Thalamic effects across clinical conditions

A main aim and strength of our study was the ability to examine multiple brain disorders jointly using a unified analytical pipeline. We found consistently smaller volumes of the medial and lateral thalamic groups in SCZ, DEM, MCI and MS. These thalamic structures overlap with the so-called *thalamic matrix*, as indicated by expression of the marker protein calbindin (Figure 3). The thalamic matrix is canonically associated with higher order cognitive functions, i.e. “higher order thalamus” (64), in contrast to the *thalamic core* (64, 65), which is mainly associated with sensory functions. While this characterization does not preclude involvement in thalamo-cortical loops within core regions such as the pulvinar nuclei, it has been found that the core and matrix regions are involved in different connectivity patterns (66, 67). Reduced medial thalamus volume may be related to the cognitive symptoms observed across ASD, SCZ, DEM, MCI, and MS, and the medial thalamic structure may thus be an important target for diagnosis or intervention. Intriguingly, we found that the thalamic core presented relatively normal volumes. The opposite effects between matrix and core regions suggest heterogeneous disease effects within the thalamus that may be related to functional characteristics (e.g. participation in cortical and subcortical circuits), local microstructural properties, or a combination of both. Further studies directly assessing the involvement of the higher-order versus sensory thalamus are needed, as our results indicate that this functional subdivision has implications for understanding the comorbidity and symptoms overlap across brain disorders.

Although overall patterns of thalamic structural alterations were similar across some of the disorders, there was some divergence between them. We found that neurodevelopmental conditions (ASD, ADHD) showed a different pattern of volumetric differences (larger volumes) relative to controls than neurodegenerative or psychiatric disorders with onset later in life (smaller volumes). Further comparisons showed that diagnoses with similar symptom profiles were preferentially clustered together (Figure 3.c), the degree of alteration differed between diagnoses (Figure 3.f), the number of affected nuclei varied (Figure 4), and the different conditions tended to distribute differently within a common components space (Figure 5). Furthermore, some conditions formed clusters in different areas of the component space, which may reflect condition heterogeneity or disease stage. These exploratory analyses suggest that thalamic nuclei volume delineate a landscape of variation across common brain disorders that can be leveraged to study comorbid diagnoses and overlapping traits (13).

A notable result is that the bilateral lateral geniculate nuclei (LGN) showed significant effects across several disorders. As part of the core thalamus, the LGN relays retinal information to primary visual cortices. Recent rodent studies suggest the LGN functions more complexly than a simple relay station, as its activity is modulated before reaching the cortex (44, 68, 69). Notably, the ventral LGN receives projections from the higher order thalamus, including the medio-dorsal thalamus (70), and is linked to circadian rhythms via connections to the suprachiasmatic nuclei (71-74). Disruptions in sleep patterns and circadian activity are common in brain disorders (75-77), warranting further research into LGN subregions’ roles in these pathophysiological conditions.

### Thalamic effects within clinical conditions

Associations between thalamic volume differences and psychiatric and neurological disorders have been reported previously. However, this is the first study to examine thalamic nuclei groups and individual nuclei across a broad range of such conditions.

We found that DEM and MCI were associated with reduced thalamic volumes across a large portion of the thalamus, showing the highest numbers of affected nuclei in the case-control comparisons, and largely overlapping patterns. Furthermore, in these diagnoses, anterior, lateral and medial thalamic groups volume were positively correlated with cognitive status as measured using MMSE. Interestingly, the volume of the medial group was also correlated with MMSE score in healthy controls, which may indicate a general association of medial thalamic size with cognitive impairments in the elderly population. Our results extend multiple previous studies reporting lower thalamic volumes in Alzheimer’s disease and MCI (22, 27, 78). Furthermore, we found overlapping patterns of DEM and MCI at the individual nuclei level, indicating possible spatially localized effects of thalamic change along the MCI-DEM continuum.

We also found widespread thalamic group and nuclei-specific atrophy in MS. A significant negative correlation between clinical function (as measured by EDSS) and volume in anterior, lateral, medial and posterior thalamic groups indicates that smaller volumes are related to more severe symptoms. On the other hand, ventral nuclei such as the ventral postero-lateral, presented larger volumes, a pattern that was similar in DEM and MCI. This suggests that somatosensory relay nuclei such as the VPL (79) may not be as susceptible in neurological diseases as compared with non-relay nuclei. The longitudinal analysis showed an overall reduction in whole thalamic volume but no thalamic group-specific effects. As a methodological consideration, the number of scans decreased for follow-up sessions. However, we repeated the longitudinal analysis only for participants with three sessions (n = 117), and the results were consistent. Reductions in thalamic volume is a well-known marker in MS (25, 80-83), as well as thalamic volume reductions over time (84, 85). Our results indicate that anterior, medial, lateral and posterior regions of the thalamus are the most affected in MS and relate to symptom severity, that posterior nuclei reduction may be a unique feature of thalamic change in MS, and that thalamic volume declines over time after disease onset.

Smaller volumes in medial and lateral groups were also found in SCZ. Interestingly, the case-control comparison at the individual nuclei level and the dimension reduction approach revealed different patterns between SCZ and the neurological disorders. Many previous studies reported smaller thalamic volume in SCZ (23, 26, 86, 87) and specifically in the medial and lateral nuclei (88), even at early stages of the disorder or in individuals at high risk. Specific thalamic alterations have been suggested to be a hallmark of schizophrenia (28, 89). Of note, many studies have reported smaller pulvinar volumes in SCZ (23, 87, 90, 91). We found significantly reduced posterior volumes when we did not correct for whole thalamic volume in the analyses (Figure 2.a), which agrees with the reported reduction in pulvinar volumes, explains differences in findings across studies, and suggests that pulvinar reductions are a large contributor to overall thalamic reductions. Furthermore, other studies observed reductions in medial geniculate nuclei, which we have not observed here (87). While this could have been the result of the correction for whole thalamic volume, other studies suggest that it is related to presence of hallucinations (91). Our results suggest that delving deeper into these effects may give greater insights into both shared and disorder-specific features along the schizophrenia spectrum.

In our large sample, we found significantly larger whole thalamic volume in ASD, as well as larger anterior thalamic group and smaller medial group volumes. A significant enlargement in ASD has also been reported in a recent study, driven by larger size in posterior regions (92). Our results suggest that this effect may be driven by larger volumes of the lateral and inferior pulvinar, as revealed in the case-control analysis at the individual nuclei level (Figure 4.a). We also found an enlargement in the anterior group in ADHD. Larger thalamus in ADHD has been related to higher symptom severity (93). Our samples covered a wide age range (from childhood to adulthood) and age specific effects may have gone undetected. The different pattern in ASD and ADHD (i.e., enlargement) compared to DEM, MCI and MS (reductions) may be related to differences between early developmental processes and later deafferentation or neurodegenerative mechanisms. In the case of SCZ, it is possible that both neurodevelopmental and degenerative processes occur, but our cross-sectional data did not allow us to disentangle these effects. Recent transcriptomic studies have identified gene expression components that are more relevant for neurodevelopmental disorders than for neurodegenerative disease (94, 95). For instance, neurodevelopmental disorders may be related to deficiencies in early basic cellular function and energetics processes in the brain (94) that may affect the establishment of thalamo-cortical circuits (96), whereas neurodegenerative disorders may involve deafferentiation and consequent synaptic dysfunction or neuronal loss (27).

We found smaller volume in BD for lateral and posterior thalamic groups. The pattern correlated with the pattern of SCZ (Figure 3.b), however the thalamic group volumes in BD were not significantly different from healthy controls after correction for whole thalamic volume. A recent study on thalamic nuclei using the same segmentation approach found smaller medial and posterior nuclei in BD (26). Our results are consistent with those findings and suggest that the effects accompany a change in whole thalamic volume.

In contrast to the other neurological conditions, we found moderately larger thalamic volume in PD. This increase was general across the thalamus and not specific to a particular nuclei group. Previous results on thalamic volume in PD are variable, in general indicating no significant volumetric differences but some shape differences between patients and controls (97-99). A recent study (N = 131 patients) reported larger thalamic volume in patients compared to controls (100). The evidence thus points towards either no volume change or a general enlargement of the thalamus as a whole in PD, without specific nuclei involvement. Of note, the patients of the current study were in the early phase of the disorder, and none used anti-Parkinson drugs. The mechanisms underlying early-phase enlargement of some brain regions in Parkinson’s disease are not fully understood, and tissue swelling due to early disease-related damage, inflammatory responses, or compensatory neuroanatomical changes are potential contributors (101, 102).

We found no significant volume differences in MDD and SCZRISK, possibly due to a small sample size or larger heterogeneity of these conditions. One possibility is that the sample sizes lacked the power to detect small differences. Another possibility is that the patients were scanned at earlier disease stages that do not yet show large changes in the thalamus. Indeed, MDD has been associated with smaller thalamic volumes (103) but only in patients who did not achieve remission (104). Although we observed this trend, it was not significant. Similarly, a recent study described smaller medial volumes in at-risk mental state individuals, although the effects were small and only significant at the most liberal threshold (88). We also found a particular reduction in the medial thalamus after correcting for whole thalamic volume, but this effect was not significant. Interestingly, MDD, SCZRISK and PSYMIX were closer in the dendrogram. Although we can only hypothesize as to why the thalamic abnormalities were less pronounced in SCZRISK and MDD, it is expected that SCZRISK presents less severe abnormalities than SCZ. Furthermore, genetic links between MDD and SCZRISK have been identified (105).

### Methodological considerations

The current study assessed case-control differences in thalamic nuclei volume in a large sample covering multiple clinical conditions using a unified analytical pipeline. All samples included more than 100 individuals in both patient and control groups, except for SCZRISK (N = 70). The large sample sizes should permit detection of moderate effect sizes, although statistical power was limited for detecting subtle differences in the smallest groups. We used a thalamus segmentation tool (52) that is currently implemented in FreeSurfer. The segmentation accuracy is known to be limited for some nuclei with weaker signal (78). However, studies comparing the segmentation results of this tool with histological delineation of the thalamus have shown that it is successful in identifying most of the nuclei (52), with particularly high reliability in medio-dorsal thalamus (contributing approximately 70% of the volume of the medial group) and LGN (contributing approximately 10% of the volume of the posterior group) (52, 106). Recent methods that improve on the available segmentation approaches by including the acquisition or estimation of additional images (107, 108), offer the opportunity to confirm our results and further characterize thalamic structures features in health and disease. Furthermore, we applied a thorough visual inspection of all datasets to ensure that high quality data was used for the analyses. To increase robustness, we pooled the individual nuclei data into six thalamic groups. We assessed the issue of multi-site data by including only patient samples for which a control sample was available from the same scanner. While RELIEF is effective for samples with large number of individuals, it may introduce some variability for sites with few individuals. This may have been the case for some of our smaller samples and the longitudinal analysis, reducing our ability to detect significant effects.

## Conclusions

By examining thalamic subdivisions we found a profile of thalamic structural abnormalities across psychiatric and neurological disorders, and that medial and lateral regions, as well as lateral geniculate nuclei, appear more vulnerable to disease. The results also highlight the importance of examining thalamic nuclei separately, since opposing effects may be masked when studying the thalamus as a whole. In addition to shared alterations, distinct patterns of thalamic structure were associated with specific disorders. This is the first study to comprehensively map thalamic structures across common brain disorders, offering insights that can guide future research in therapeutic strategies.

## Acknowledgements

This work was performed on the TSD (Tjenester for Sensitive Data) facilities, owned by the University of Oslo, operated and developed by the TSD service group at the University of Oslo, IT-Department (USIT). Dr Buitelaar has been supported by the EU-AIMS (European Autism Interventions) and AIMS-2-TRIALS programmes which receive support from Innovative Medicines Initiative Joint Undertaking Grant No. 115300 and 777394, the resources of which are composed of financial contributions from the European Union’s FP7 and Horizon2020 Programmes, and from the European Federation of Pharmaceutical Industries and Associations (EFPIA) companies’ in-kind contributions, and AUTISM SPEAKS, Autistica and SFARI; and by the Horizon2020 supported programme CANDY Grant No. 847818). The European Research Council under the European Union’s Horizon 2020 research and Innovation program (ERC StG Grant No. 802998). Research Council of Norway (#324499, #324252, #344121) and Nordforsk (#164218). The funders had no role in the design of the study; in the collection, analyses, or interpretation of data; in the writing of the manuscript, or in the decision to publish the results. Any views expressed are those of the author(s) and not necessarily those of the funders. Data used in preparation of this article were obtained from the Alzheimer’s Disease Neuroimaging Initiative (ADNI) database (adni.loni.usc.edu). As such, the investigators within the ADNI contributed to the design and implementation of ADNI and/or provided data but did not participate in analysis or writing of this report. A complete listing of ADNI investigators can be found at: http://adni.loni.usc.edu/wp-content/uploads/how_to_apply/ADNI_Acknowledgement_List.pdf. Data used in the preparation of this article were obtained from the Alzheimer’s Disease Neuroimaging Initiative (ADNI) database (adni.loni.usc.edu), PPMI [http://www.ppmi-info.org/], the ABIDE project [https://fcon_1000.projects.nitrc.org/], ADHD200 [http://fcon_1000.projects.nitrc.org/], OASIS [http://www.oasis-brains.org/], SCHIZCONNECT [http://schizconnect.org/] and OpenNeuro (https://openneuro.org/). A detailed overview of the included cohorts and acknowledgement of their respective funding sources and cohort-specific details are provided in Supplementary Table 1.

